# Impaired subgranular zone radial glia morphology and transient amplification of neural progenitors in *Mllt11*-deficient mice leads to increased hippocampal neurogenesis

**DOI:** 10.1101/2025.10.02.680006

**Authors:** Sam Moore, Danielle Stanton-Turcotte, Karolynn Hsu, Emily A. Witt, Angelo Iulianella

## Abstract

The mammalian hippocampus derives from the cortical hem region of the forebrain and is one of the two regions in the mammalian forebrain that contains neural progenitors capable of generating new neurons in the postnatal brain. Hippocampal neural stem cells, referred to as type I progenitors, are Sox2^+^/GFAP^+^, localize along the hilar boundary region, and possess a radial glia fiber that spans the thickness of the dentate gurus. The hilar boundary region serves as a niche for the neural progenitor pool, controlling the activation of type-1 progenitors to transit amplifying type-2 neural progenitors which lose their apical adherence to the hilus as they differentiate to granule cells. The genetic regulation of this process is not fully understood and requires factors that control the mobilization of type-1 cells from their hilar niche in the subgranular zone. To that end we now report a role for Mllt11 (Af1q/Tcf7c; Myeloid/lymphoid or mixed-lineage leukemia; translocated to chromosome 11/All1 Fused Gene from Chromosome 1q/T cell factor 7 co-factor), a regulator of neurite formation and migration in the cortex, in the regulation of hippocampal neurogenesis by restricting the transition of type-1 cells to transit amplifiers and neuroblasts. We previously showed that Cux2 (Cutl2) is widely expressed in the developing hippocampus, initially in the subventricular zone of the cortical hem, which generates the dentate gyrus, then in both type-1 and type-2 neural progenitors, and later in maturing granule cells of the perinatal hippocampus. To evaluate a role for *Mllt11* in hippocampal neurogenesis, we used a *Cux2^IRESCre/+^* line to generate a *Mllt11* loss-of-function in perinatal hippocampal progenitors and differentiating granule cells. The loss of *Mllt11* led to enlarged dentate blades dues to increased generation of NeuN^+^ and Calbindin^+^ granule cells. Explorations of the progenitor pool revealed increased transit amplifying cells due to an expanded pool of displaced Sox2^+^/GFAP^+^ deep in the dentate blades, with altered radial glial morphology.

Consequently, type-1 progenitors aberrantly transitioned to amplifying progenitors and neuroblasts, resulting in enhanced granule cell neurogenesis in the postnatal hippocampus. Primary neurosphere formation assays confirmed enhanced proliferation of neural stem cells derived from *Mllt11* knockout hippocampi. Taken together, the findings reported here demonstrate the critical role of Mllt11 in maintaining the hippocampal radial glial phenotype and their association within the subgranular zone-hilar niche boundary region, thereby controlling their differentiation to transit amplifiers and granule cells.

## Introduction

Neurogenesis in the mammalian CNS was traditionally accepted to occur only during embryonic stages. However, it has more recently become understood that new neurons are added in discrete regions of the adult mammalian CNS, specifically the subventricular zone (SVZ) of the rostral migratory stream and the subgranular zone (SGZ) of the hippocampal dentate gyrus (DG) (Ehninger and Kempermann, 2008; Kempermann et al., 2015; Ming and Song, 2011). The hippocampus is a critical structure of the limbic system, with a principal role in memory formation, especially the transformation of short-term memory to long-term memory (Borzello et al., 2023; Hainmueller and Bartos, 2020; Sosa and Giocomo, 2021; Zeithamova and Bowman, 2020). As such, adult hippocampal neurogenesis has become an area of great interest owing to the potential role in neurodegenerative disease, cognitive decline, and neurodevelopmental disorders (Banker et al., 2021; Boekhoorn et al., 2006; Culig et al., 2022; Curtis et al., 2003; Fedele et al., 2011; Hoglinger et al., 2004; Jin et al., 2004; Lazarov and Marr, 2010; Marxreiter et al., 2013; van den Berge et al., 2011).

Development of the DG begins around E12.5 in a complex process that involves the migration of proliferative progenitors away from the dentate neuroepithelium (DNE), the primary matrix located between the hippocampal neuroepithelium and the cortical hem (Guerout et al., 2014; Moore and Iulianella, 2021; Urban and Guillemot, 2014). From a developmental point of view, this is a unique process, as the formation of fated DNE precursors migrate inward, away from their sites of origin, to generate an SVZ of proliferating cells. At E14.5, cortical hem-derived Cajal-Retzius (CR) cells secrete Reelin, a glycoprotein, to aid the migration of these precursors toward the pial side of the cortex to form the secondary germinal matrix from which the hippocampal dentate neuroepithelium arises (Hodge et al., 2013; Seri et al., 2004; Sugiyama et al., 2013; Urban and Guillemot, 2014; Yamada et al., 2015). At this time, radial glial precursors begin to form hippocampal neurons in the adjacent hippocampal neuroepithelium. By E17.5, dentate precursors accumulate in the newly established hippocampal fissure to form the tertiary matrix (Sugiyama et al., 2013; Yamada et al., 2015). As hippocampal neurons are born, they migrate along radial glial cells toward their location in the hippocampal cornu ammonis (CA) fields. During DG development, precursors from all three matrices produce granule cells that form the outer-granule cell layer (GCL) (Sugiyama et al., 2013; Urban and Guillemot, 2014). The DG ultimately forms in an outside-in manner, as the continued proliferation of immature granule cells and their upward migration contributes to the inner-GCL (Mathews et al., 2010; Yamada et al., 2015). At birth, the characteristic blade shape of the DG forms, dictated by CR cells. GC neurons in the DG first appear below the hippocampal fissure, to establish what is known as the upper blade of the DG. While continuous migration of CR cells promotes the formation of the lower blade (Amaral et al., 2007). Precursors in the primary and secondary matrix soon disappear, restricting proliferation to the tertiary matrix or SGZ during postnatal DG development (Alvarez-Buylla and Lim, 2004; Sugiyama et al., 2013; Urban and Guillemot, 2014).

Progenitors within the hippocampus transition through a series of cell types, ending in the formation of the mature functionally integrated granule cell neurons (Ambrogini et al., 2004; Kempermann et al., 2015; Pleasure et al., 2000; Zhao et al., 2006). Specifically, type-1 radial glia-like neural progenitors (NPs) rapidly give rise to intermediate neuronal progenitors, with type-1 (radial glia), type-2a (transit amplifier), and type-2b (neuroblast) cells maintaining the potential for self-renewal and amplification (Ambrogini et al., 2004; Kempermann et al., 2015; Pleasure et al., 2000; Roybon et al., 2009; Seri et al., 2004; Steiner et al., 2006). Type-3 cells exit the cell cycle to mature into newborn neurons, ultimately integrating into existing neural networks as mature granule cells. Between P0 and P14, most hippocampal granule cell neurons are born and begin consolidating to form axonal connections targeting neurons in the CA1 region of the hippocampus (Yasuda et al., 2011). The early postnatal period is also critical for the generation of the adult neural progenitor pool, with neurogenesis declining over time due to a loss of radial glial type I neural progenitors (Ben Abdallah et al., 2010; Encinas et al., 2011; Encinas and Sierra, 2012; Yamada et al., 2015).

Here we report a role for Mllt11/Af1q (Myeloid/lymphoid or mixed-lineage leukemia; translocated to chromosome 11/All1 Fused Gene from Chromosome 1q) in regulating hippocampal neurogenesis. Mllt11 is a novel vertebrate-specific protein that was initially defined as an oncogenic factor fused to Mll to create an aberrant product identified in acute myeloid leukemia patients carrying the t(1;11)(q21;q23) translocation (Tse et al., 1995). However, the role of the unfused protein, referred to Mllt11 (or Af1q, Tcf7c) during normal development was unknown. Work from our laboratory has begun characterizing Mllt11 as a novel regulator of neural development. *Mllt11* is highly and exclusively expressed in post-mitotic neurons during development and enriched in the developing cortical plate (Yamada et al., 2015). We previously demonstrated that *Mllt11* is required to promote nascent neuron migration and neuritogenesis in the cortex (Stanton-Turcotte et al., 2022), retina (Blommers et al., 2023), cerebellar granule cells (Blommers et al., 2024), and progenitors of the lateral ventricle choroid plexus (Moore et al., 2025). We previously described *Mllt11* mRNA expression in the postnatal hippocampus, which was lost in *Mllt11* cKOs (Stanton-Turcotte et al., 2022), but the role of Mllt11 in hippocampal development and neurogenesis is unknown.

To address this, we used a *Cux2^IRESCre/+^* driver line to inactivate *Mllt11* in the developing dentate germinal epithelium and progenitors lining the SGZ using an *Mllt11^flox/flox^* mouse line.

Cux2 is expressed in neural progenitors in the SGZ from embryonic day (E)14.5 onwards, and later in type-1 and type-2 postnatal hippocampal progenitors, with a bias towards Sox2^+^/GFAP^-^ type-2 activated neural progenitors in the first two weeks of life, declining thereafter (Yamada et al., 2015). This reflects the activity of Cux2 in an activated subset of hippocampal neurogentic progenitors during the peak neurogenic phase of early postnatal life. We showed through lineage tracing using the *Cux2^IRESCre/+^* mouse strain that Cux2 activity not only labeled neurogenic progenitors in the early postnatal hippocampus but also marks the formation of GCL neurons in an outside-in manner (Yamada et al., 2015), making it a suitable Cre driver line to inactivate *Mllt11* during hippocampal neurogenesis. We now report that *Mllt11* loss enhanced neurogenesis in the mammalian hippocampus, primarily due to enhanced transition of type-1 neural progenitors to type-2 transit amplifiers. *Mllt11* cKO hippocampi displayed reduced radial glial morphology, which led to an aberrant basal translocation of the Sox2^+^/GFAP^+^ neural progenitor population within the developing DG. This was associated with the *Mllt11* mutant hippocampal progenitors rapidly transitioned to amplifying neural progenitors and neuroblasts, and enhanced formation of granule cells in the DG. Taken together, the findings described here demonstrate the critical role of Mllt11 in maintaining the hippocampal radial glial phenotype and the balance between neural progenitor proliferation and terminal differentiation.

## Materials and Methods

### Animals

Mice used in this study were handled in accordance with the regulations of Dalhousie animal ethics committee and the guidelines of the Canadian Council on Animal Care.

*Mllt11^flox/flox^; Rosa26^tdTomato/TdTomato^*conditional targeted reporter allele were generated as described previously (Stanton-Turcotte et al., 2022). To inactivate *Mllt11^flox/flox^* and activate *Rosa26^tdTomato^* in the developing dentate gyrus and hippocampal progenitors, a *Cux2^IRESCre/+^*driver line used, which we previously characterized (Stanton-Turcotte et al., 2022; Yamada et al., 2015). The *Cux2^IRESCre/+^; Ai9 (Rosa26^tdTomato/TdTomato^)* mouse line targets *TdTomato* transgene activation initially *to* the SVZ of the cortical *hem*, which is fated to give rise to the DG and associated progenitor cells, and later granule cell neurons, reflecting an outside-in maturation gradient. *Cux2^IRESCre/+^* was maintained in a C57Bl6/J background and crossed into a tdTomato reporter line maintained in FVB background to generate *Cux2^IRESCre/+^; Rosa26^tdTomato/tdTomato^*. This line was then crossed with *Mllt11^flox/flox^; Rosa26^tdTomato/tdTomato^* to generate *Cux2^IRESCre/+^; _Rosa26_tdTomato/tdTomato_; Mllt11_flox/+* _to be backcrossed with *Mllt11*_*flox/flox_; Rosa26_tdTomato/tdTomato*_. All_ experiments were performed with *Cux2^IRESCre/+^; Rosa26^tdTomato/tdTomato^; Mllt11^+/+^* as the wild type Cre^+^ control (WT), and *Cux2^IRESCre/+^; Rosa26^tdTomato/tdTomato^; Mllt11^flox/flox^* as the conditional knockout or cKO, as we previously described (Stanton-Turcotte et al., 2022).

### Histology and immunohistochemistry

Whole brains were dissected out at embryonic day 18.5 (E18.5) and postnatal stages (P)7, P14 and 6 weeks after birth. For postnatal stages litters were anesthetized by intraperitoneal injection of 100μl of 4:1 Ketamine:Xylazine followed by Perfusion with PBS and 4% PFA. Whole brains were fixed in 4% PFA (0.1m phosphate buffer) at 4 for 2 hours to overnight depending on the embryonic or postnatal stage. This was followed by 3 x 10-minute washes in PBS and cryoprotection in 15%-30% sucrose. Once equilibrated, whole brains were embedded and snap frozen in Optimum Cutting Temperature compound (O.C.T.; Tissue-Tek, Torrance, CA) and stored at -80 . A total of 40 coronal sections per animal at 12µm per section were obtained from WT and cKO. Immunohistochemistry was conducted as previously described (Iulianella et al., 2008). The following primary antibodies were used: rabbit anti-Calbindin (1:1000, Abcam), rabbit anti-Pax6 (1:500, Hybridoma), goat anti-Sox2 (1:200, Santa Cruz), mouse anti-NeuN (1:200, Millipore), goat anti-NeuroD1 (1:300, Santa Cruz), rabbit anti-Tbr1 (1:200, Millipore), goat anti-Nestin (1:200, Santa Cruz), rabbit anti-Prox1 (1:1000, Sigma), mouse anti-Calretinin (1:500), rabbit anti-Cleaved Caspase3 (1:500), rabbit anti-Mllt11 (1:50), and rabbit anti-GFAP (1:500, Abcam). Secondary antibodies were used at 1:1500 and included donkey anti-rabbit Alexa-Fluor 647, donkey anti-rabbit Alexa-Fluor 488, donkey anti-goat Alexa-Fluor 647, and donkey anti-mouse 647 (Invitrogen). Images were captured using a Zeiss AxioObserver inverted fluorescent microscope with a 20X objective and a 40X oil objective and Apotome2 Structural Illumination. Montages were assembled using Photoshop CS6 (Adobe, San Jose, CA) or Pixelmator Pro.

### In situ hybridization

30µm frozen sections were obtained from brains of WT control and cKO mice staged at P7, P14, P21, and P28. Tissue was fixed overnight as previously described, and Mllt11 riboprobe hybridization was performed as previously described (Yamada et al., 2014).

### EdU (5-ethynyl-2′-deoxyuridine) in vivo labeling

10mM EdU solution was prepared in DMSO and administered via intraperitoneal injection into dams. EdU was dosed at 30mg/kg body in a solution of 10mg/ml PBS (pH 7.35). A series of time points for injection and harvesting were used for full assessment of hippocampal neurogenesis. The dosing time points were as follows: dose at E14.5, harvest at E18.5; dose at E16.5, harvest at E18.5; and finally dose at E18.5, harvest at P14. EdU staining was conducted using Click-iT EdU kit according to the manufacturer’s protocol (Invitrogen). The immunohistochemistry protocol was adapted such that EdU staining was performed before the addition of the primary antibody, as described previously (Blommers et al., 2023).

### Sampling methodology

To ensure consistency among samples, cell counts obtained at E18.5, P7, P14 and 6 weeks after birth, were restricted to a total of 10 counting frames (50μm x 50μm) randomly placed on the entire DG starting from the midline to the lateral end of the DG according to systematic-random sampling method. The base of each counting frame was adjusted to align along the lower edge of the SGZ to ensure cells within the hilus region were not included in counts. A total of 40 coronal sections were obtained from 3-4 separate individuals at 12μm thickness obtained from WT or cKO mice at E18.5, P7, P14 and 6 weeks. To analyze DG thickness, equivalent coronal sections were selected and examined from each *WT* control and conditional *Mllt11* cKO mutant animal and repeated for a total of 10 mice. Measured values were averaged to create one representative value for each animal. A portion of the DG with uniform thickness was selected as a rectangular area of interest. The area was then divided by the length of the rectangular area with the thickness calculated from the proximal areas to the crest where the two blades connect.

### Hippocampal neurosphere formation assays

Primary and secondary neurosphere growth assays were performed as previously described (Fatt et al., 2015). Briefly, P7 hippocampi from WT control and Mllt11 cKO littermates were dissected in Hanks’ Buffered Saline Solution (HBSS) supplemented with antibiotics (Penicillin-Streptomysin, Invitrogen), and cells were dissociated with a Trypsin/Hyalurondidase/Kurenic acid enzyme mix (Sigma-Aldridge) in HBSS at 37°C for 30 minutes on a shaker. Trypsin Inhibitor (Sigma-Aldridge) was added to the mix and cells were triturated and spun down at 1500rpm for 5 minutes. Cells were resuspended in Serum Free medium (DMEM low glucose-F12, 30% glucose, 7.5% NaHCO3, 1M HEPESpH7.4, L-glutamine, Penicillin-Streptomycin; Invitrogen) supplemented with FGF2 (StemCell Technologies), EGF (StemCell Technologies), Heparin (StemCell Technologues), and B27 (Invitrogen). Cell counts were performed using a Nile Blue (Invitrogen) exclusion assay and plated at 10 cells/μL and cultured for 1 week and counted at the end of the culture period. For secondary sphere assays, cells were spun down and resuspend as above in fresh serum-free media, passed through a 10μm cell strainer, counted using the Nile Blue exclusion assay, and plated at 2 cells/μL. Cells were cultured for 1 week, after which they were counted.

### Statistics

All statistical analyses were performed using PRISM (Graphpad). Statistical significance was obtained by performing unpaired t-tests to compare the means and standard deviations between the control and the experimental data sets. In all quantification studies, statistical differences were challenged using the Student’s t-test (two-tailed), with significance level set at P ≤ 0.05 (*P ≤ 0.05, **P ≤ 0.01, ***P ≤ 0.001, ****P ≤ 0.0001). For all experiments, a minimum of 3 WT control and 3 cKO individuals were analyzed and cell counts were reported as the mean of the values for each individual (n identifies the number of individuals per each WT and cKO data set). For neurosphere assays, cells were derived from microdissected hippocampi from a total of 7 cKO and 3 WT Cre^+^ controls at P7. Cell counts were performed blinded to genotype to avoid potential counting bias. Data was presented as scatter boxplots, with each point representing an individual and results presented as mean ± Standard Error of the Mean (SEM).

## Results

We previously described the neuronally-restricted *Mllt11* expression in the developing nervous system, including the forebrain (Stanton-Turcotte et al., 2022; Yamada et al., 2014). Using a conditional loss-of-function approach we showed that *Mllt11* is required for the migration of superficial cortical neurons, neuritogenesis and formation of callosal projections (Stanton-Turcotte et al., 2022), the migration of newborn neurons in the retina (Blommers et al., 2023) and cerebellar granule cells (Blommers et al., 2024), and migration of cortical hem neuroepthelial progenitors cells that contribute to the formation of the lateral ventricle choroid plexus (Moore et al., 2025). However, the role of Mllt11 in hippocampal development, which also arises from the cortical hem region (Moore and Iulianella, 2021), is unknown. To address this, we first characterized the expression of *Mllt11* in the postnatal hippocampus by *in situ* hybridization (ISH) in C57BL/6 mice over a series of postnatal time points. At postnatal day P7 and P14 *Mllt11* mRNA was intensely expressed in the DG and CA1-CA3 regions, with expression declining at P21 and P28, suggesting a role in perinatal neurogenesis (Figure S1A-D). We next utilized a Cre/LoxP strategy to generate *Mllt1l* cKO using a *Cux2^IRESCre/+^*driver mouse strain, along with a tdTomato reporter allele to monitor targeted cells. *Cux2* is dynamically expressed in the developing mouse forebrain, cortical hem, and dentate gyrus (Yamada et al., 2015; Zimmer et al., 2004). This makes the *Cux2^IRESCre/+^* driver a suitable choice to inactivate *Mllt11* during hippocampal neurogenesis. We examined the DNE at E14.5 and the DG at E18.5 and noted largely normal hippocampal morphology in the fetal brains of *Mllt11* cKOs, except for a shortened choroid plexus (Fig. S1E-H), which was profiled phenotypically in a recent publication (Moore et al., 2025). Lineage tracing of tdTomato^+^ cells in *Cux2^IRESCre/+^; Rosa26r^tdTomato/tdTomato^* brains from E18.5, P7 and P14 confirmed Cre activity in the developing DNE, DG, and choroid plexus (Fig. S1I-N) (Yamada et al., 2015). We previously reported that the *Cux2* locus is active in the developing fetal DNE at E14.5 and E18.5, which includes neural progenitors seeding the DG (Yamada et al., 2015). At postnatal stages, *Cux2^IRESCre/+^*activity becomes progressively restricted to nascent GCs, reflecting the maturation of the DG in an outside-in manner (Fig. S1K-N) (Yamada et al., 2015). The developing DG appeared shorter in *Mllt11* cKOs relative to controls at E18.5 (Fig. S1I, J), while at postnatal stages tdTomato^+^ staining was expanded in the DG of *Mllt11* cKOs at P7 and P14 relative to controls, and the dentate blades exhibited increased thickness (Fig. S1K-N). This suggested that *Mllt11* loss led to enhanced postnatal hippocampal neurogenesis. We therefore examined dentate morphometrics using DAPI staining and found that dentate blade thickness was indeed increased in *Mllt11* cKO relative to controls, particularly at perinatal stages at P7 (Fig. S2A-C, J; p<0.0001, n=10) and P14 (Fig. S2D-F, J; p<0.0001, n=10). A slight but significant increase in dentate blade thickness was also observed in*Mllt11* cKOs at 6 weeks of age, when mice are sexually mature (Fig. S2G-I, J; p< 0.05, n=3). We probed for changes in apoptosis by staining for the presence of Cleaved Caspase-3^+^ cells and observed no change between *Mllt11* cKOs and controls at E18.5, P7, or P14 (Fig. S2K, L; E18.5: p=0.52, n=4; P7: p=0.28, n=4; P14: p=0.11, n=4).

To explore this further we investigated the generation of mature granule cells using the nuclear phosphoprotein maker NeuN and the calcium-binding protein Calbindin, which labels migratory immature granule cells in the dentate gyrus (Marques-Mari et al., 2007; Yoo et al., 2011). NeuN cell counts displayed a surprising decrease in *Mllt11* cKO relative to controls at E18.5 (Fig. 1A-C; p<0.05, n=3), in contrast with a significant increase in NeuN^+^ nuclei in the DG of *Mllt11* cKOs at postnatal stages P7 (Fig. 1D-F; p<0.01, n=3) and P14 (Fig. 1G-I; p<0.05, n=3). Interestingly, we observed an increase in tdTomato^+^ fate labeling of the GCL of *Mllt11* cKO at P7, indicating enhanced granule cell maturation and neurogenesis at perinatal stages (Fig.1J-M). We observed a similar pattern of early reduction of fetal granule cell neurogenesis upon *Mllt11* loss at E18.5 (Fig. S3A-C; p<0.01, n=4), with a contrasting increase in Prox1^+^ cells at P7 (Fig. 6; p=0.13, n=4) and P14 (Fig. 6; p<0.05, n=4), demonstrating enhanced GCL neurogenesis. This is consistent with *Cux2^IRESCre/+^* tdTomato^+^ fate-labeling, which showed a truncated DNE in cKOs at E18.5 (Fig. S1K, L), reflecting the activity of the Cre driver line from E14.5 onwards in the cortical hem and SVZ contributing to DNE development. Given that our *Cux2^IRESCre/+^* tdTomato^+^ fate-labeling studies demonstrated enhanced granule cell neurogenesis following *Mllt11* loss (Fig. S1I-N), we next characterized the nascent neuronal pool by profiling Calbindin staining in the DG. As with NeuN (Fig. 1), the conditional deletion of *Mllt11* resulted in a transient decrease in the number of Calbindin^+^ cells at E18.5 (Fig. S4A-C; p<0.05, n=3) and at P7 (Fig. S3D-F; p<0.01, n=3), but no significant difference was reported at P14 (Fig. S3G-I; p=0.8, n=3). In higher magnification views, the expansion in Calbindin^+^ cells emerging from the SGZ of the *Mllt11* cKO DG was clearly visible (Fig. S3J, L) and was accompanied by expansion in *Cux2^IRESCre/+^*-driven tdTomato^+^ fate-labeled cells (Fig. S3K, M), reflecting increased granule cell neurogenesis.

**Figure 1.**
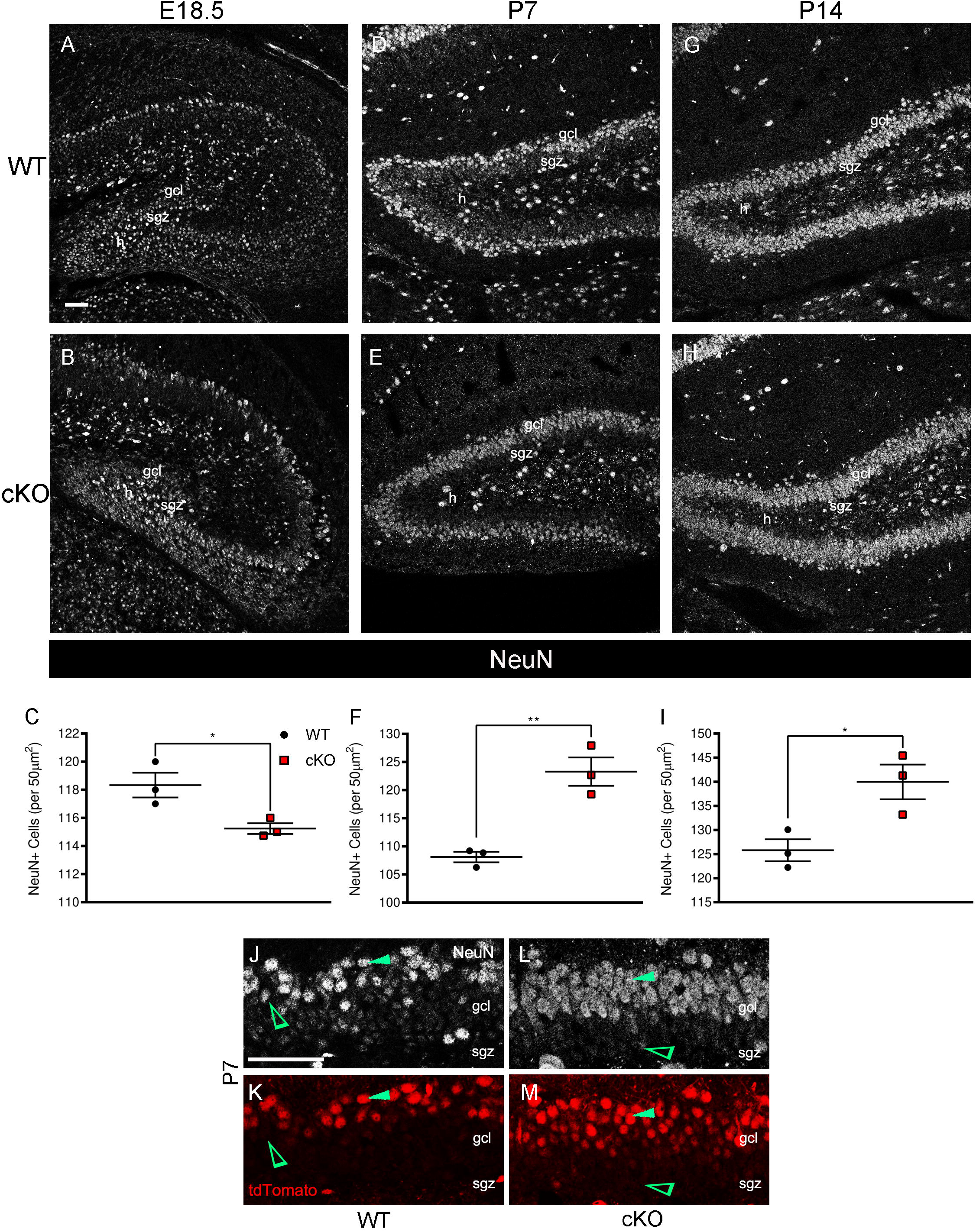
*Mllt11* loss increased formation of NeuN^+^ mature granule cells in the postnatal dentate gyrus. (A, B) Coronal sections of the developing DK at E18.5 in the WT and conditional *Mllt11* cKO mouse stained for NeuN (white) identifying mature GCs. (C) Scatter boxplot identifying a significant decrease in mature GCs in the cKO relative to WT controls (P< 0.05, n=3). (D, E) Coronal sections of the dorsal hippocampus at P7 in the WT and cKO mouse stained for NeuN. (F) Scatter boxplot identifying a significant increase in mature GCs in the cKOs (P< 0.01, n=3). (G, H) Coronal sections of the dorsal hippocampus at P14 in the WT and cKO mouse stained for NeuN. (I) Scatter boxplot identifying a significant increase in mature GCs in the cKOs (P<0.05, n=3). (J-M) Magnified cross-sectional view of the DG at P7 identifying the SGZ and GCL of WT and cKO. Top panel: NeuN staining; lower panel: recombined tdTomato^+^ cells. (J-M) Solid green arrowhead identifying NeuN^+^/tdTomato^+^ mature GC. Open green arrowhead identifying NeuN^+^ mature GC with no tdTomato expression. Scale bars = 50µm. Data presented as mean ± SEM. Abbreviations: dk, dentate knot; gcl, granule cell layer; h, hilus; sgz, subgranular zone.

*Mllt11* loss led to aberrant maturation of hippocampal granule cells within the first two weeks of life, however, when examining the findings from Calbindin and NeuN staining more closely, a complex role of *Mllt11* in promoting the maturation of GCL neurons emerges. Our previous work identified biphasic Cux2 activity during hippocampus development. From E14.5-16.5 Cux2 is active in the hippocampal forming region of the SVZ in the CH region of the medial forebrain, while at postnatal stages Cux2 is expressed in an activated cohort of type-1 progenitors as they are transitioning to type-2 amplifying cells, as well as in maturing DG granule cells (Yamada et al., 2015). Thus, the effect of *Mllt11* loss in the maturation of cells in the DG may reflect the progressive deletion of the floxed *Mllt11* allele in progenitors and GCL neurons when using the *Cux2^IRESCre/+^* mouse driver. To more clearly delineate the effect of *Mllt11* loss on hippocampal neurogenesis, we evaluated the formation of nascent Calbindin^+^ GCL neurons using a series of EdU birth dating studies. Three EdU pulse-chase time points were chosen to best reflect the outside-in formation of the DG, as we previously reported (Yamada et al., 2015). Pulsing with EdU at E14.5 and harvesting at E18.5 was conducted to capture the generation of earliest granule neurons from the primary germinative matrix residing in the SVZ of the CH. Pulsing with EdU followed by harvesting at E16.5 and harvesting at E18.5 was intended to capture the formation of the earliest dentate blade, which is truncated in *Mllt11* cKO mutants (Fig. S1I, J). The third EdU experiment involved pulsing at E18.5 and harvesting at P14 to capture the switch to nascent GCL production from within DG after its formation, reflecting postnatal neurogenesis. The conditional deletion of *Mllt11* in the hippocampal primordium led to a significant increase in of EdU^+^/Calbindin^+^ cells at E18.5 when pulsed at E14.5 (Fig. 2A-C, J-M, p< 0.001; WT: n=3, cKO: n=4). This suggested that *Mllt11* loss led to a transient spike of GCL neurogenesis, as these EdU^+^/Calbindin^+^ cells comprise the earliest born GCL neurons populating the outer edge of the DG blade (Yamada et al., 2015). In contrast, EdU pulsing at E16.5 revealed a significant decrease in the cohort of Calbindin^+^/EdU^+^ double positive cells in cKO neonates at E18.5 (Fig. 2D-F, p<0.05, n=4), confirming a transient delay in dentate blade formation in the *Mllt11* cKOs, as revealed by tdTomato^+^ fate mapping (Fig. S1K, L). In contrast, EdU pulse labeling at E18.5 and analysis at two weeks postnatally (P14), confirmed that conditional *Mllt11* loss enhanced neurogenesis of the earliest-born GCL neurons (Fig. 2 G-I, N-Q, p<0.01, n=3). When taken together with the DG post-mitotic marker analysis, the EdU pulse-labeling experiment confirmed a role for Mllt11 in regulating the pace and extent of hippocampal neurogenesis. Specifically, *Mllt11* loss initially attenuated the contribution of granule cell neurons progenitors from the dentate knot as they migrate to fill the blades of the DG, but once the dentate blades are formed, *Mllt11* loss led to a burst GCL neurogenesis.

**Figure 2.**
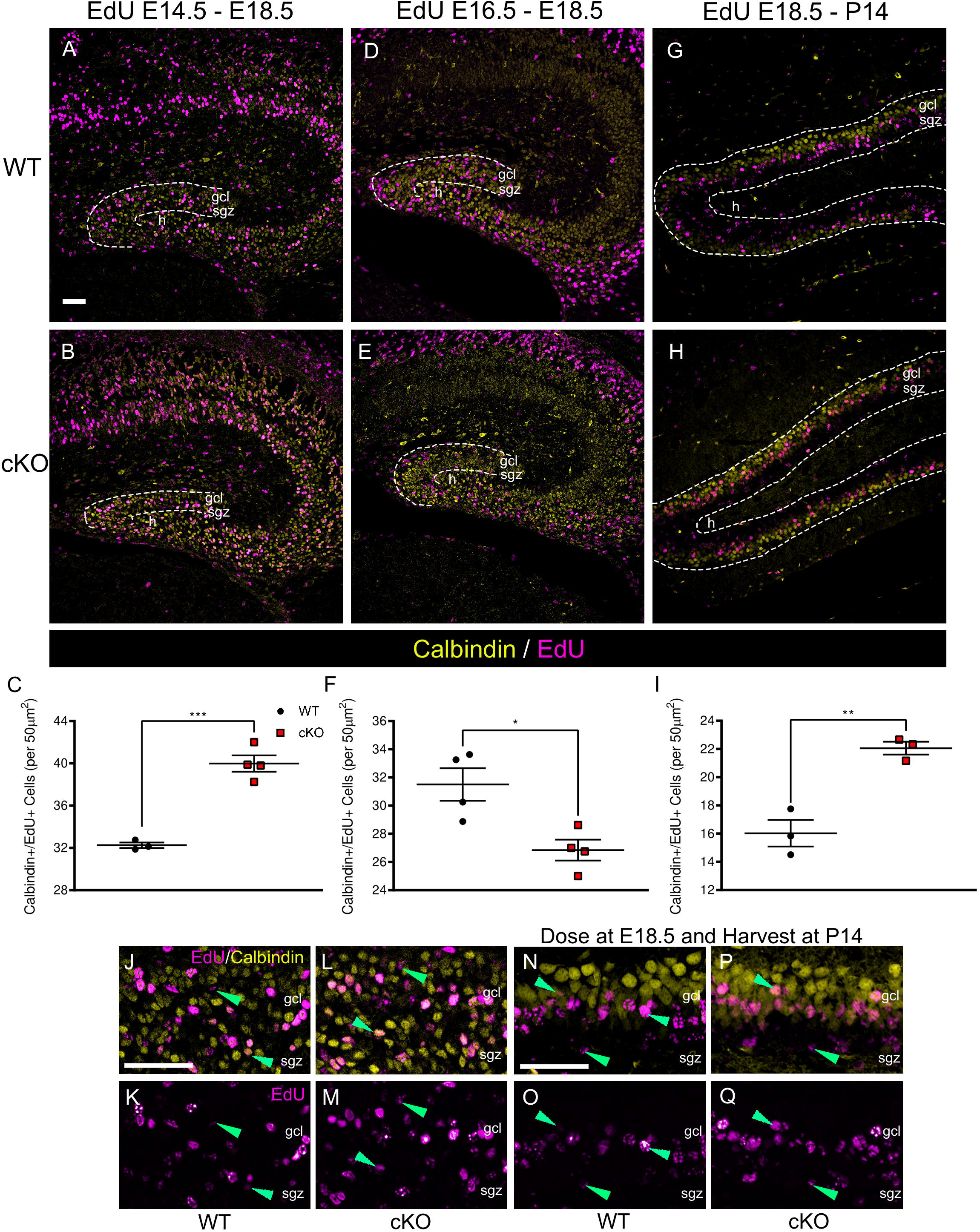
*Mllt11* loss enhanced postnatal neurogenesis of granule cells. (A, B) Coronal sections of the developing DK in the WT and *Mllt11* cKO mouse dosed with EdU at E14.5 and harvested at E18.5, stained for Calbindin (yellow) and EdU (violet). (C) Scatter boxplot quantifying a significant increase in Calbindin^+^/EdU^+^ cells in the cKO in comparison to WT controls (P<0.001, WT: n=3, cKO: n=4). (D, E) WT and cKO E18.5 coronal sections across the developing hippocampus pulsed with EdU at E16.5 and stained for Calbindin and EdU. (F) Scatter boxplot identifying a significant decrease in Calbindin^+^/EdU^+^ cells in cKOs (P< 0.05, n=4). (G, H) Coronal sections of the dorsal hippocampus at P14 in the WT and cKO mouse stained for Calbindin and EdU. (I) Scatter boxplot quantifying a significant increase in Calbindin^+^/EdU^+^ cells in the cKOs (P< 0.01, n=3). (J-M) Magnified cross-sectional view of the DG of WT and cKO mice dosed with EdU at E14.5 and harvested at E18.5, identifying labeled cells in the SGZ and GCL. Top panel: Calbindin^+^/EdU^+^ cells; lower panel: EdU^+^ labeling. (J-Q) Magnified cross-sectional view of the DG of WT and cKO mice dosed with EdU at E18.5 and harvested at P14, identifying the SGZ and GCL. (J-Q) Top panel: overlay of Calbindin and EdU channels; bottom panel: EdU^+^ cells. Green arrowhead identifying Calbindin^+^/EdU^+^ cells. Scale bars = 50µm. Quantification of data presented as mean ± SEM. Abbreviations: dk, dentate knot; gcl, granule cell layer; h, hilus; sgz, subgranular zone.

To delineate the cellular mechanism underlying the transient increase in perinatal neurogenesis following *Mllt11* ablation, we next profiled the progenitor pools of the SGZ using Pax6, and Sox2/GFAP co-staining to identify type-1 progenitors within the SGZ exhibiting radial glial morphology, and Sox2^+^/GFAP^-^ and NeuroD1^+^ to identify type-2a transient amplifiers and type-2b neuroblasts, respectively (Roybon et al., 2009; Steiner et al., 2006). Type-1 hippocampal progenitors originate at fetal stages from the SVZ region of medial telencephalon known as the dentate knot (DK), and express the radial glial progenitor markers GFAP, Sox2, and Pax6 (Catalani et al., 2002; Ehninger and Kempermann, 2008; Kempermann et al., 2015; Komitova and Eriksson, 2004; Steiner et al., 2006; Suh et al., 2007). These progenitors comprise a subset of astroglia which go on to populate the SGZ of the hilar niche bordering the dentate blade, possess radial processes that span the thickness of the dentate blade, and generate newborn GCs postnatally (Kempermann et al., 2015; Seri et al., 2004). Type-2 cells represent a developmental state in which the proliferative transient amplifying (type-2a) cells quickly differentiate into neuroblasts (type-2b) and then to immature neurons (type-3 cells) in the DG.

We first examined the distribution of Pax6^+^ cells at E18.5, P7, and P14, encompassing the period of perinatal neurogenesis, and noted an initial modest increase in the number of Pax6^+^ cells per unit area at E18.5 (Fig. 3A-C; p<0.01, n=4), followed by a decrease at P7 (Fig. 3D-F, J-K; p=0.0001, n=5) and no significant change at P14 (Fig. 3G-I; p=0.42, n=5). Thus, *Mllt11* loss may have caused a transient burst of neural progenitor proliferation followed by neurogenesis (Fig. 1,2, S4) and reduction of the progenitor pool at perinatal stages. To directly test for this possibility, we undertook a neurosphere formation assay from control (*Cux2^IRESCre/+^*) and *Mllt11* cKO (*Cux2^IRESCre/+^; Mllt11^flox/flox^*) P7 hippocampi, as described previously (Fatt et al., 2015). We noted that *Mllt11* loss enhanced primary neurosphere formation (Fig. 3L: p<0.001, WT: n=3, cKO: n=7) but did not affect secondary neurospheres proliferation (Fig. 3M; p=0.35, WT: n=3, cKO: n=7). While this is consistent with the observed burst of GCL neurogenesis we observed in the cKOs, the lack of an effect on secondary neurosphere formation was puzzling. Primary neurospheres are derived from dissociated neural progenitors from the hilar niche, whereas secondary neurospheres derive from the isolated daughter progenitors generated by the primary progenitor pool, and as such lack contact with the hilus, suggesting that *Mllt11* loss affected neural progenitor interactions within the hilar niche.

**Figure 3.**
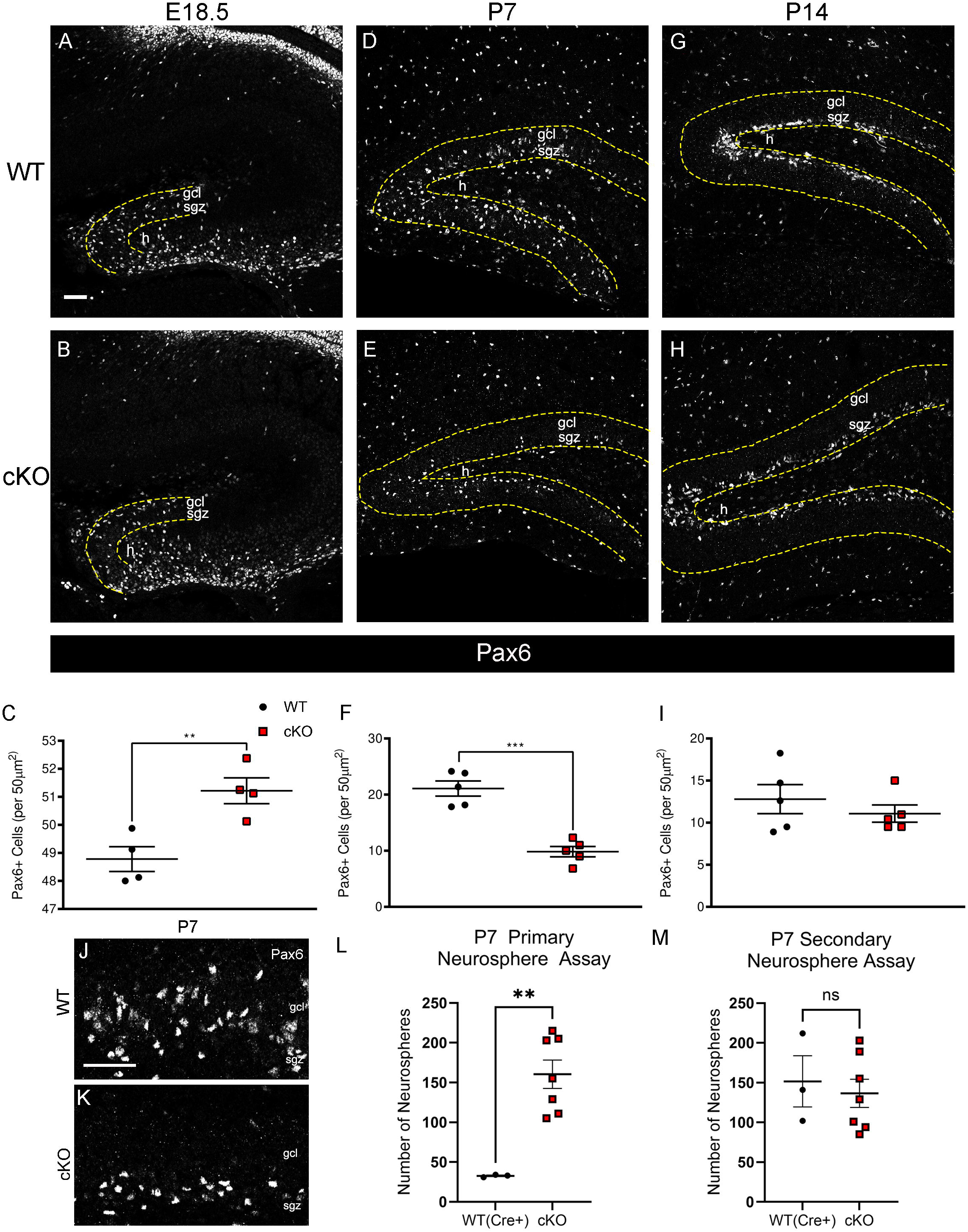
*Mllt11* loss leads to a transient expansion and then exhaustion of perinatal hippocampal progenitors. (A, B) Coronal sections of the developing DK at E18.5 in the WT and *Mllt11* cKO mouse stained for Pax6 (white) identifying type-1 hippocampal progenitors. (C) Scatter boxplot identifying a significant difference in type-1 cells in the cKO in comparison to WT controls (P<0.01, n=4). (D, E) Coronal sections of the dorsal hippocampus at P7 in the WT and cKO mouse stained for Pax6. (F) Scatter boxplot identifying a significant decrease in type-1 cells in the cKO relative to WT controls (P<0.0001, n=5). (G, H) Coronal sections of the dorsal hippocampus at P14 in the WT and cKO mouse stained for Pax6. (I) Scatter boxplot quantifying no significant difference in type-1 cells in cKOs compared to WT controls (P= 0.42, n=5). (J, K) Magnified cross-sectional view of the DG identifying the SGZ and GCL of WT and cKO stained for Pax6. (L, M) Quantification of primary (L) and secondary (M) neurosphere formation from isolated P7 hippocampal neural progenitors from WT (*Cux2^IRESCre/+^*) and cKO (*Cux2^IRESCre/+^; Mllt11^flox/flox^*). Mllt11 cKOs exhibited increased primary neurosphere production (L) (p<0.001, WT: n=3, cKO: n=7) but no change in secondary neurosphere formation. (M) (p=0.35, WT: n=3, cKO: n=7). Scale bars = 50µm. Data presented as mean ± SEM. Abbreviations: dk, dentate knot; gcl, granule cell layer; h, hilus; sgz, subgranular zone.

To explore this further we profiled the morphology of the radial glial pool of hippocampal progenitors identified by GFAP^+^/Sox2^+^ co-staining during perinatal development. We also examined Sox2^+^/GFAP^-^ type-2a cells in the SGZ, which reflect the activated pool of neural progenitors. As with the Pax6 data, *Mllt11* cKO mutants showed an initial increase in the GFAP^+^/Sox2^+^ type-1 radial glial progenitor pool at E18.5 (Fig. 4A-C; p<0.01, n=3), and an increase in type-2a Sox2^+^/GFAP^-^ cells (Fig. 4C; p<0.01, n=3). In contrast at P7 and P14, *Mllt11* loss led to a reduction in the number of co-stained GFAP^+^/Sox2^+^ type-1 radial glial progenitors (P7: Fig. 4D-F, p<0.001, n=5; P14: Fig.4G-I, p<0.001, n=4), and a corresponding transient increase in the number of Sox2^+^/GFAP^-^ type-2a cells at P7 (Fig. 4F; p=0.0001, n=5), but not at P14 (Fig. 4G-I; p=0.14, n=4). Interestingly, many of the Sox2^+^ cells were unmoored from the hilus and instead displaced basally from the SGZ and into the dentate blade in P7 *Mllt11* cKO mutants, with their radial glial processes failing to span the thickness of the DG (green open arrowheads, Fig. 4J, K). This suggested that *Mllt11* loss altered the stability of radial glial process of type-1 neural progenitors and consequently led to their expansion and enhanced transition to type-2a amplifiers within the first week of life, accounting for the observed burst in GCL neurogenesis.

**Figure 4.**
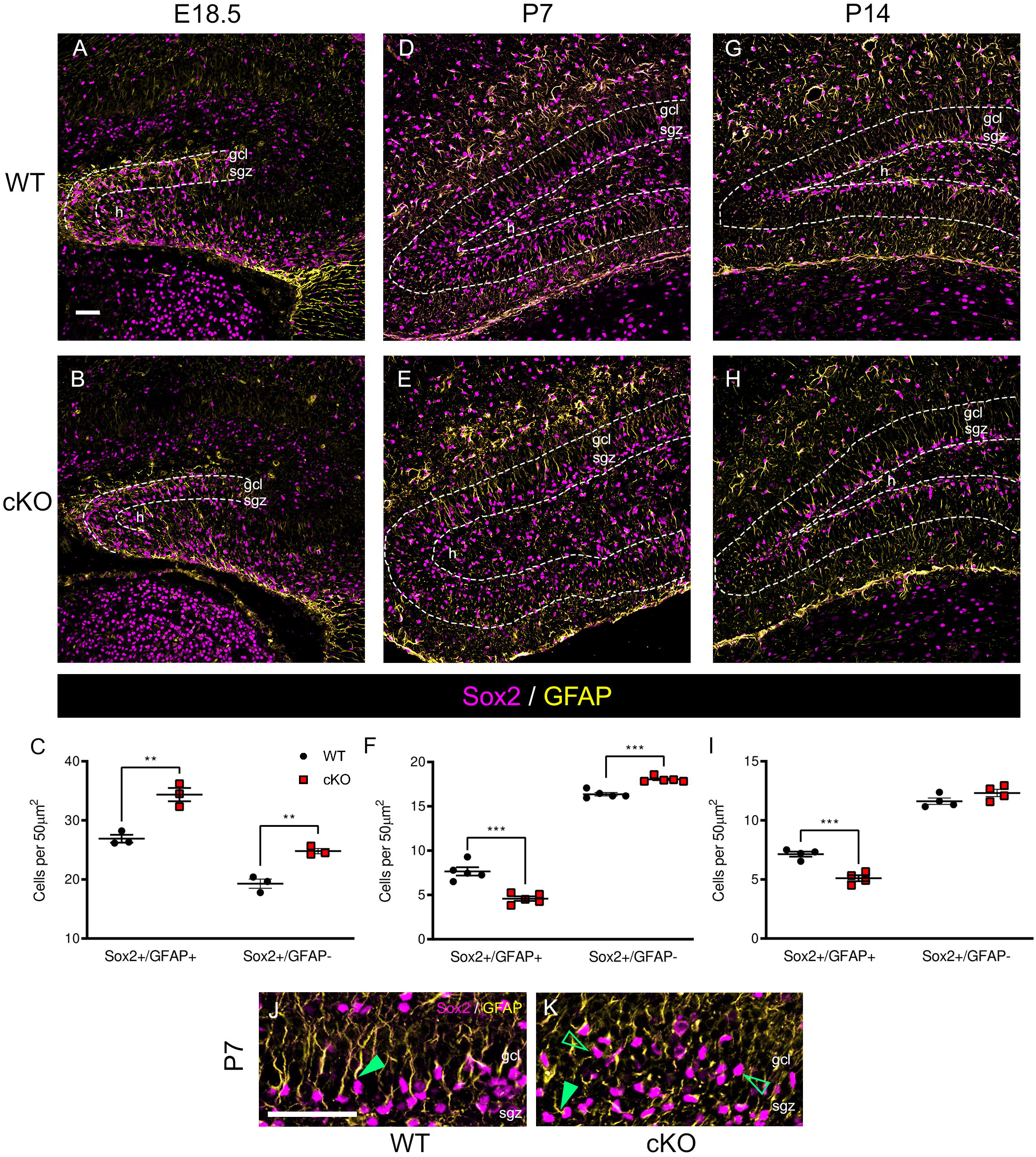
*Mllt11* loss impacted the stability of radial glial processes and anchoring of type-1 cells to the hilar niche. (A, B) Coronal sections of the developing DK at E18.5 in the WT control and *Mllt11* cKO mouse brain stained for Sox2 (violet) and GFAP (yellow). Sox2^+^/GFAP^+^ co-stain identifies type- 1 cells and Sox2^+^/GFAP^-^ cells identifies type-2a cells. (C) Scatter boxplot identifying a significant increase in both type-1 (p<0.01, n=3) and type-2a (p< 0.01, n=3) cells in the embryonic DK region in cKOs compared to WT controls. (D, E) Coronal sections of the dorsal hippocampus at P7 in the WT and cKO mouse stained for Sox2 and GFAP. (F) Scatter boxplot identifying a significant decrease in type-1 cells (p<0.001, n=5) with a corresponding increase in type-2a cells (p= 0.0001, n=5). (G, H) Coronal sections of the dorsal hippocampus at P14 in the WT and cKO mouse stained for Sox2 and GFAP. (I) Scatter boxplot identifying significant decrease in type-1 cells (p<0.001, n=4) with no significant difference in type-2 cell numbers (P= 0.14, n=4) in cKO sections compared to WT controls. (J-K) Magnified cross-sectional view of the DG identifying the SGZ and GCL of WT and cKO stained for Sox2 (violet) and GFAP (yellow). Solid green arrowheads in (J-K) identify GFAP^+^/Sox2^+^ type-1 NP cells. Open green arrowheads in (J-K) identify type-2a Sox2^+^/GFAP^-^ NPs. (J, K) P7 *Mllt11* cKO displayed reduced GFAP^+^ radial glial fibers, basally displaced Sox2^+^ cells from the SGZ, and expanded Sox2^+^/GFAP^-^ cells. Scale bars = 50µm. Data presented as mean ± SEM. Abbreviations: dk, dentate knot; gcl, granule cell layer; h, hilus; sgz, subgranular zone.

Given that *Mllt11* loss expanded the Sox2^+^/GFAP^-^ neural progenitors in the DG, we expected to see changes in the pool of migrating granule cell neuroblasts in the postnatal hippocampus. Neuroblasts, or type-2b cells, are progenitors that have left their niche and begun to terminally differentiate into granule cell neurons. They are characterized by expression of Cux2 (Yamada et al., 2015), and NeuroD1, a pro-neural transcription factor (Roybon et al., 2009; Steiner et al., 2006). *Mllt11* loss led an increase in NeuroD1^+^ cells in the DG relative to WT control at all time points observed (Fig. 5A-I; E18.5: p=0.001, n=4; P7: p=0.0001, n=3; P14: p=0.002, n=4), consistent with a increase in Sox2^+^/GFAP^-^ type-2a transit amplifying cells (Fig.4A-I). NeuroD1-staining confirmed a slightly truncated DG region at E18.5 in *Mllt11* cKOs relative to controls (Fig.5 A, B) but showed a significant increase in NeuroD1^+^ cells upon quantification of cell numbers (Fig. 5C). Altered DG formation was also observed with markers for GCs (Figs. 1, S3, S4) and radial glial progenitors (Figs. 3, 4). Interestingly, there was a pronounced increase in NeuroD1^+^ cells in the hilus region of the *Mllt11* cKO hippocampus, which was most notable at P7 (not included in the cell count data; Fig. 5D, E). This suggested that the expansion of radial glial progenitors in the *Mllt11* cKO hippocampus led to a quick burst of highly migratory perinatal neuroblast formation within the first week of life. This is sufficient to account for the enlarged DG blades (Fig. S2) and postmitotic GC in the *Mllt11* cKO brains (Fig. 1, S3, S4). The increase in NeuroD1^+^ type-2b neuroblast formation was still observed in *Mllt11* cKOs at P14, reflecting increased perinatal neurogenesis (Fig. 5G-I). This coincided with expanded Pax6^+^ and Sox2^+^/GFAP^+^ type-1 NPs in cKOs at P7 (Figs. 3, 4). Altogether, our ontogenetic analysis demonstrated that *Mllt11* loss impacted the maintenance of hippocampal progenitors most acutely within the first week of life, leading to a greatly expanded population of transit amplifiers.

**Figure 5.**
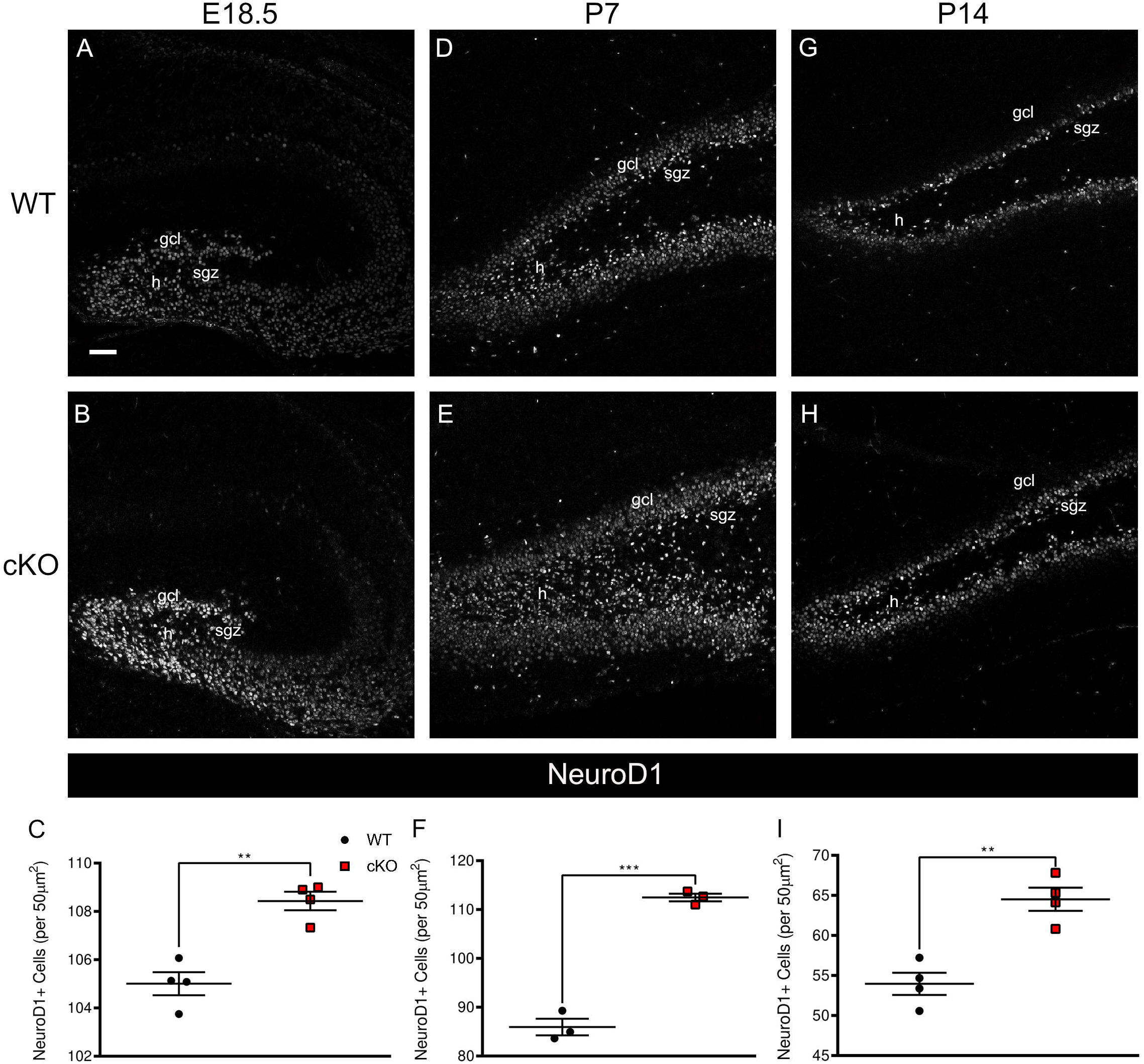
Enhanced NeuroD1^+^ type-2b transit amplifiers in the hippocampus following *Mllt11* inactivation. (A, B) Coronal sections of the developing dk at E18.5 in the WT control and *Mllt11* cKO mouse stained for NeuroD1 identifying type-2b transit amplifying cells (neuroblasts). (C) Scatter boxplot identifying a significant increase in NeuroD1^+^ neuroblasts in the cKOs relative to WT controls (P= 0.001, n=4). (D, E) Coronal sections of the dorsal hippocampus at P7 in WT and cKO DGs labeling NeuroD1. (F) Scatter boxplot quantifying a significant increase in neuroblasts in the cKOs compared to WT controls (P= 0.0001, n=3). (G, H) Coronal sections of P14 WT and cKO DGs stained for NeuroD1. (I) Scatter boxplot identifying a significant increase in neuroblasts in the cKOs compared to WTs (P= 0.002, n=4). Scale bars = 50µm. Data presented as mean ± SEM. Abbreviations: dk, dentate knot; gcl, granule cell layer; h, hilus; sgz, subgranular zone.

Next, we explored the impact of *Mllt11* loss the terminal phase of hippocampal DG neurogenesis. The stage is characterized by the maturation of granule cells, whose dendrites branch towards the molecular layer of the DG, and axons project towards the hippocampal CA3 pyramidal cell layer. Newly generated GCL neurons become postmitotic and are identified by markers for immature granule cells, including Prox1 (Lavado et al., 2010), a prospero type homeodomain protein, and Tbr1, a T-box transcription factor (Englund et al., 2005; Hevner et al., 2006). The loss of *Mllt11* resulted in significantly fewer Prox1^+^ cells being generated at E18.5 (Fig. 6A-C; p<0.01, n=4). In contrast, there was a slight trend toward an increase in Prox- 1^+^ cells at P7, which was not statistically significant (Fig. 6D-F; p=0.14, n=4), and a significant increase at P14 (Fig. 6G-H; p<0.05, n=4). Interestingly, no differences were noted in the numbers of Tbr1^+^ cells at E18.5 (Fig. 7A-C; p=0.4511, n=3), but significant increases were found in the DG of *Mllt11* cKOs at postnatal stages; namely P7 (Fig. 7D-F; p<0.01, n=3) and P14 (Fig. 7G-I; p=0.01, n=4). High magnification views of the DG at P7 confirmed that Tbr1^+^ nuclei expressed no (open green arrowheads) or low (filled green arrowheads) levels of *Cux2^IRESCre/+^; tdTomato^+^* labeling (Fig. 7J-M). High levels of tdTomato^+^ transgene expression were restricted to the outermost GCL, reflecting Cux2 activity during the maturation of postmitotic neurons, which were expanded in *Mllt11* cKOs compared to controls (Fig. 7J-M).

**Figure 6.**
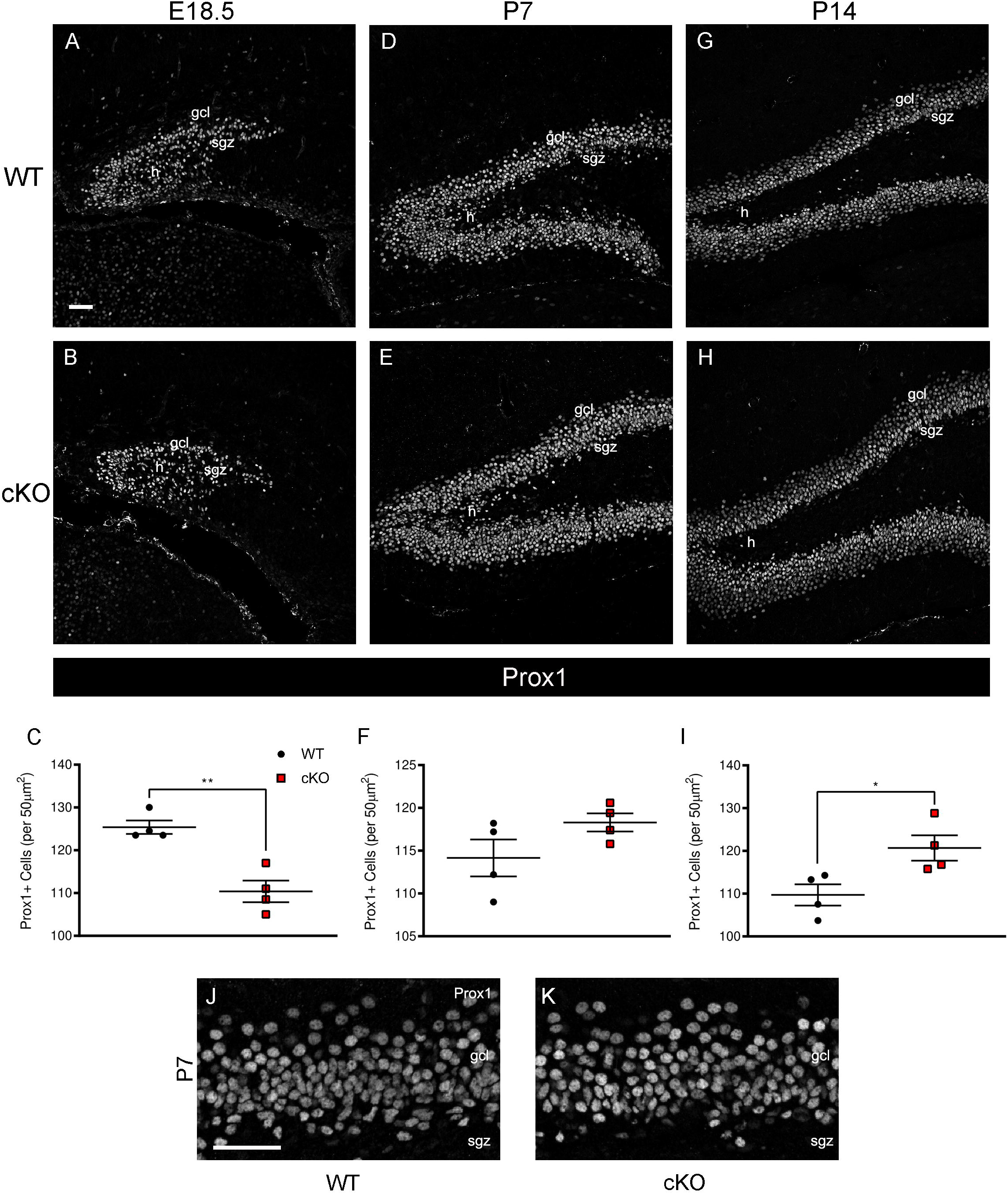
*Mllt11* loss altered the formation of Prox1^+^ granule cells in the developing dentate gyrus. (A, B) Coronal section of the developing DK at E18.5 in the WT control and *Mllt11* cKO mouse stained for Prox1, identifying maturing GCs. (C) Scatter boxplot quantifying a significant decrease in maturing GCs in the cKOs at E18.5 (P< 0.01, n=4). (D, E) Coronal sections of the dorsal hippocampus at P7 in the WT and cKO mouse stained for Prox1. (F) Scatter boxplot identifying non-significant increase in maturing GCs in the cKOs relative to WT controls at P7 (P= 0.13, n=4). (G, H) Coronal sections of the dorsal hippocampus at P14 in the WT and cKO mouse stained for Prox1. (I) Scatter boxplot identifying a significant increase in maturing GCs in the cKOs vs. WTs at P14 (P< 0.05, n=4). (J, K) Magnified cross-sectional view of the DG at P7 identifying the SGZ and GCL of WT and cKO stained for Prox1. Scale bars = 50µm. Data presented as mean ± SEM. Abbreviations: dk, dentate knot; gcl, granule cell layer; h, hilus; sgz, subgranular zone.

**Figure 7.**
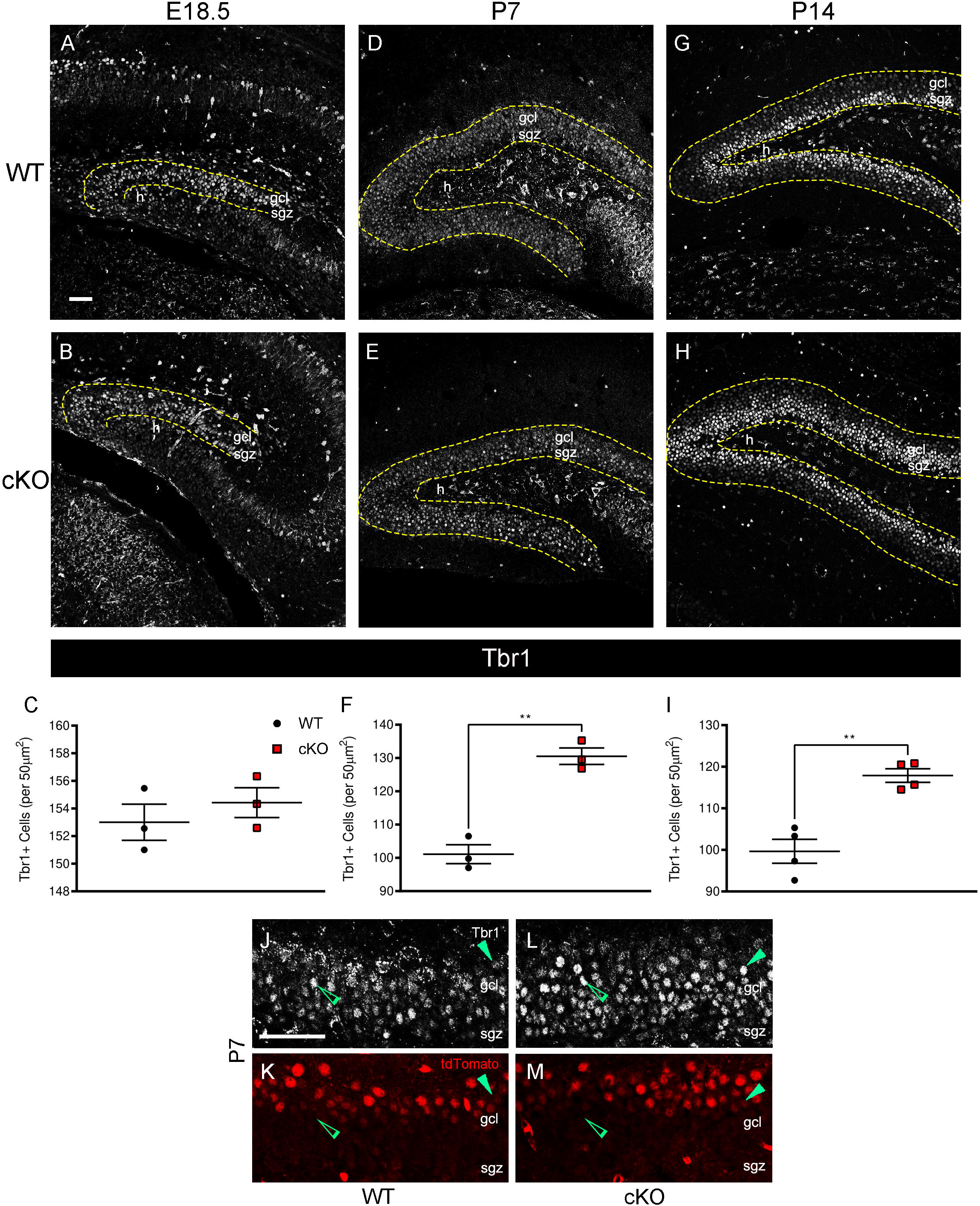
*Mllt11* loss increased formation of Tbr1-positive developing granule cells. (A, B) Coronal sections of the developing DK at E18.5 in the WT and *Mllt11* cKO mouse stained for Tbr1 (white) identifying maturing GCs. (C) Scatter boxplot identifying no significant difference in maturing GCs in the cKO in comparison to WT controls (P= 0.45, n=3). (D, E) Coronal sections of the dorsal hippocampus at P7 in the WT and cKO mouse stained for Tbr1. (F) Scatter boxplot quantifying a significant increase in maturing GCs in the cKOs (P< 0.01, n=3). (G, H) Coronal sections of the dorsal hippocampus at P14 in the WT and cKO mouse stained for Tbr1. (I) Scatter boxplot quantifying a significant increase in maturing GCs in the cKOs compared to WT controls (P<0.01, n=4). (J-M) Magnified cross-sectional view of the DG at P7 identifying the SGZ and GCL of WT and cKO; top panel: Tbr1 stain; lower panel: recombined tdTomato^+^ cells. (J-M) Solid green arrowheads identify maturing GC with high Tbr1 intensity and low tdTomato levels. Open green arrowhead identifying maturing GC with low Tbr1 intensity and high tdTomato levels. Scale bars = 50µm. Data presented as mean ± SEM. Abbreviations: dk, dentate knot; gcl, granule cell layer; h, hilus; sgz, subgranular zone.

Taken together, these findings demonstrate that *Mllt11* loss initially impacted the maintenance of type-1 NPs at perinatal stages, with cKO mutant cells displaying altered radial glial morphology with reduced adherence to the hilar nice. This then led to a rapid transition of type-1 cells to type-2 transit amplifying progenitors within the first week of life, ultimately contributing to the enhanced maturation of the GCL within the first two weeks of life, resulting in increases in expression of markers for immature (Prox1, Tbr1) and mature (Calbindin, NeuN) GCs.

## Discussion

The objective of this study was to characterize the role of *Mllt11* in hippocampal development and neurogenesis. We previous reported on the generation of a *Mllt11* loss-of-function conditional mouse mutant which exhibited neuronal migration defects in the cortex (Stanton-Turcotte et al., 2022), retina (Blommers et al., 2023), cerebellum (Blommers et al., 2024) and choroid plexus (Moore et al., 2025). Yet its role in the hippocampus was unknown. The hippocampus is one of two regions of the mammalian brain that can undergo neurogenesis postnatally, with the bulk of neurogenesis occurring within the first two weeks of life (Yamada et al., 2015). Of critical importance to the genesis of new neurons is the DG neural niche, a region straddling the interface between the dentate blade and hilus regions, populated by a collection of radial glial neural progenitors (type-1 cells) and activated transit amplifiers (type-2a cells) (Duan et al., 2008; Kempermann et al., 2015; Roybon et al., 2009; Urban and Guillemot, 2014; Yamada et al., 2015). The maintenance of the radial glia morphology and hilar niche interactions are thought to be essential for a lifelong maintenance of neurogenic capacity of the hippocampus.

We now report a role for Mllt11 in regulating hippocampal neurogenesis primarily by maintaining the radial glial morphology of hippocampal progenitors, which is essential for their adherence to the hilar niche region. In the absence of *Mllt11*, there was a loss of Sox2^+^/GFAP^+^ type I cells and corresponding expansion of Sox2^+^/GFAP^-^ and NeuroD1^+^ type-2 transit amplifying cells, which led to enhanced granule cell production in the postnatal hippocampus. Using neurosphere formation assays we also showed that *Mllt11* loss stimulated proliferation of primary neural progenitors in culture, further corroborating that its loss led to the aberrant activation of neural progenitors to transit amplifiers.

Embryonic and postnatal neurogenesis relies on the unique spatiotemporal sequence of cell type differentiation to maintain a necessary balance between the generation of hippocampal progenitors and terminal differentiation yielding mature postmitotic neurons. We previously showed that the *Cux2^IRESCre/+^* mouse strain activates *tdTomato* reporter gene within the developing dentate neuroepithelium of the SVZ region in the cortical hem (Yamada et al., 2015). In the first week of life, Cux2 is activated in two patterns consistent with hippocampal neurogenesis: (1) it is expressed in some type-1 cells but mostly in the Sox2^+^ and GFAP^-^ (type- 2a) activated neural progenitor pool residing within the SGZ, and (2) it is activated in the DG in an outside-to-inside pattern, reflecting the maturation of the GCL (Moore and Iulianella, 2021; Yamada et al., 2015). The *Cux2^IRESCre/+^*strain is therefore well-suited for targeting the inactivation of a *Mllt11^flox/flox^*allele in the perinatal hippocampus, including neural progenitors undergoing the transition to amplifiers. Using this genetic loss-of-function strategy, we observed that *Mllt11* acts by maintaining the radial glial architecture of type-1 cells which is required for their positioning along the SGZ niche. Proper radial glial morphology is essential for the long-term self-renewal of hippocampal progenitors (Falk and Gotz, 2017; Steiner et al., 2006). In the absence of *Mllt11*, type-1 cells appeared untethered, and their glial processes are shortened and fail to maintain a connection from the hilus/SGZ boundary to the top of the DG blade. We suggest that the reduction of hilar-niche interaction of *Mllt11* mutant hippocampal progenitors, led to transient increases in the numbers of amplifying progenitor cell types, such as NeuroD1^+^ type-2 and Tbr1^+^ immature GCL neuroblasts, concomitant with a burst in the formation of Prox1^+^ and Calbindin^+^ immature granule cells in the DG. The DG of *Mllt11* cKOs were also thicker as early as 1 week of age, as they harbored more NeuN^+^ GCL neurons. The increased hippocampal neurogenesis was also evidenced by the expanded tdTomato^+^ fate-mapped cells in the DG of the *Cux2^IRESCre/+^-*driven *Mllt11* cKOs, which showed a dramatic increase in the infilling of the DG blades in the perinatal *Mllt11* mutant hippocampus. This effect was most notable at P7, with *Mllt11* cKO mutants showing increases in the number of tdTomato^+^ cells fated to the GC identity, suggesting enhanced activation of GC fate. This was likely a consequence of the transient burst of intermediate progenitors observed in the mutant hippocampus. The increased proliferation of intermediate progenitors and terminal GC differentiation in the cKO brains was supported by EdU birth dating analysis, which showed that *Mllt11* loss resulted in a significant increase in the numbers of Calbindin^+^/EdU^+^ cells.

Our findings demonstrate a primary role of Mllt11 in the maintenance of a hippocampal radial glial phenotype and that, in its absence, cells rapidly transition through a series of developmental states ultimately leading to enhanced GC neurogenesis coupled with a reduction of self-renewing progenitor populations. The balance between the generation of hippocampal NPs and terminally differentiated postmitotic neurons shifts during development, with postnatal and adult NPs becoming molecularly and anatomically distinguishable from their embryonic counterparts (Ehninger and Kempermann, 2008; Hardwick and Philpott, 2014; Kempermann et al., 2015; Urban and Guillemot, 2014). Specifically, embryonic NPs are characterized by their high proliferative rate within a dynamic niche environment in which they reside, while adult NPs are characterized by their acquisition of quiescence within the complex but stable cellular niche at the hilus-SGZ border region (Kempermann et al., 2015; Urban and Guillemot, 2014). This shift during development is essential to the formation of the appropriately sized population of neurons and must be tightly controlled. We demonstrated that the balance between hippocampal progenitors and formation of postmitotic neurons is controlled by Mllt11 during development. In the absence of *Mllt11*, we observed an increase in “unmoored” activated NPs, accompanied by an expansion of intermediate progenitors. This implies a shift in the maintenance of NP proliferation and commitment to the neuronal cell fate due to exhaustion of the type-1 radial glial NP pool. Thus, at late fetal to perinatal stages, Mllt11 controls a shift in the balance of progenitor cell activity toward self-renewal vs. differentiation. In support of this, we showed that *Mllt11* loss was accompanied by enhanced proliferation of primary neural progenitors in primary neurosphere assays.

Mllt11 is a 90 amino acid vertebrate-specific protein whose structure is poorly defined, confounding predictions regarding interacting proteins. However, we recently identified components of the cytoskeleton as possible Mllt11 interacting proteins in the developing brain (Blommers et al., 2024; Stanton-Turcotte et al., 2022). Particularly, we conducted a proteomic screen using E18.5 brain lysates and identified tubulins, non-muscle myosins, including non-muscle myosin heavy chain 2A/non-muscle Myosin IIA (NMIIA/Myh9), Non-muscle myosin heavy chain 2B/Non-muscle Myosin IIB (NMIIB/Myh10), as potential Mllt11-interacting proteins (Blommers et al., 2024; Stanton-Turcotte et al., 2022). This suggests that Mllt11 may be a novel cytoskeletal-associating protein in brain tissues, consistent with its role in promoting neuronal migration and neuritogenesis. Interestingly, NMIIA functions in the translocation of the actin cytoskeleton and is one of three affected genes to be implicated in autism, schizophrenia and intellectual disability (Li et al., 2016). Non-muscle myosin heavy chain 2B (NMIIB) is also an actin-associating motor protein whose disruption leads to cancer metastasis, hydrocephalus, autism and schizophrenia, and is implicated in neuronal migration and neuritogenesis (Betapudi et al., 2006; Ma et al., 2007; Ma et al., 2009; O’Roak et al., 2012; Wang et al., 2018). While the roles of NMIIA and NMIIB in NP development have yet to be elucidated, we propose that they play a role in the formation and/or maintenance of the radial glial phenotype in conjunction with Mllt11. This would then explain the loss of the radial glial phenotype arising from *Mllt11* loss in the developing DG. Future studies are exploring the role of Mllt11-NMIIA/B interactions in radial glial formation of the mammalian hippocampus.

In conclusion, our study identifies Mllt11 as a key regulator of the type-1 radial glia phenotype and transition to amplifying NPs in the perinatal hippocampus. Mice lacking *Mllt11* displayed enhanced proliferation of neural progenitors and generation of GC neuroblasts, ultimately producing increased granule cell neurogenesis. The effect of *Mllt11* loss is most likely a consequence of abnormal radial glia morphology of the type-1 cells in the mutants. Our fundings support the notion that proper radial glial morphology and niche interactions are crucial for regulating the tempo of hippocampal neurogenesis and preventing the exhaustion of NPs.

However, Mllt11 may also play a role in regulating the expression of differentiation/survival factors required for cell cycle exit and terminal differentiation of GCL neurons. Future work should explore the role of the Mllt11 interactome in the maintenance of hippocampal radial glial morphology. In addition, it would be interesting to explore behavioural consequences of *Mllt11* loss using the *Cux2^IRESCre/+^* driven conditional cKOs. Tests assaying spatial memory, such as the Morris water maze (Vorhees and Williams, 2006) and the elevated plus maze (Walf and Frye, 2007) would test memory dysfunction that may arise as a result of the aberrant hippocampal neurogenesis phenotype observed in the *Mllt11* cKOs. This would provide valuable insight into the functional consequences of the inactivation of *Mllt11* during postnatal neurogenesis.

## Acknowledgements

We gratefully acknowledge funding from the National Science and Engineering Research Council of Canada (RGPIN 03925-20) and Canadian Institutes of Health Research (CIHR PJT- 388914). SM was supported by a CIHR MSc scholarship, and EW was supported by postdoctoral fellowships from the Killam Foundation and American Association for Anatomy. We thank Sarah Whitehead for assistance with animal husbandry.

## Supplementary Data Figure Legends

**Figure S1. *Mllt11* expression profiling and *Cux2^IRESCre/+^* fate mapping in the embryonic and adult hippocampus.**

(A-D) Images showing *in situ* hybridization in coronal tissue sections of C57BL/6 mice at P0, P14, P21 and P28. *Mllt11* mRNA expression detected at high levels in the DG and CA regions of the hippocampus with highest levels during peak neurogenesis at P0, dissipating by P28. Scale bar = 50µm. (E-F) DAPI staining in the DNE at E14.5 in the WT and *Mllt11* cKO mouse. Note the truncated structure of the ChP in the cKO. Scale bar = 100µm. (G-H) DAPI staining of a dorsal DG from an E18.5 cKO mouse. Scale bar = 100μm. (I-P) Coronal sections of forebrains from WT vs cKO embryos showing tdTomato (red) fluorescence as a readout for recombination in *Cux2*-expressing cells in the developing hippocampus. Top panel: coronal section from WT control mouse; bottom panel: coronal section from a comparative cKO mouse. (I, J) E18.5: *Cux2^IRESCre/+^-*mediated *tdTomato* reporter recombination in the developing gcl. (K, L) P7: highest tdTomato^+^ labeling in the gcl. (M, N) P14 tdTomato^+^ labeling in the gcl, with the occasional labeled cells in the hillus, but very few labeled cells in the sgz. Scale bar = 50µm. Abbreviations: CH, cortical hem; ChP, choroid plexus; gcl, granule cell layer; lv, lateral ventricle; sgz, subgranular zone; h, hilus.

**Figure S2. *Mllt11* loss results in an increase in dentate gyrus thickness.**

(A, B) DAPI stained coronal sections of dorsal DG at P7 in the WT and conditional cKO *Mllt11* mouse. Yellow box in (A) identifying representative region used for thickness calculation. (C) Scatter plot identifying a significant increase in DG thickness (μm) in cKO sections compared to WT controls (P<0.0001; n=10). (D, E) DAPI stained coronal sections of the dorsal DG at P14 in WT and cKOs. (F) Scatter boxplot identifying a significant increase in DG thickness (μm) in cKO sections compared to controls (P<0.0001; n=10). (G, H) DAPI stained coronal sections of the dorsal DG at 6 weeks. (I) Scatter boxplot identifying a significant increase in DG thickness (μm) in cKOs relative to WTs (P<0.05; n=3). (J) Scatter boxplot showing no significant changes in total number of DAPI^+^ cells per 50μm^2^ at E18.5 (P = 0.50; n=10), P7 (P = 0.64; n=10) and P14 (P = 0.13; n=10). (K) WT and cKO DG sections at E18.5, P7, and P14 stained for cleaved caspase-3 (yellow) and counterstained with DAPI (blue) to assess cell death. Red arrowheads indicate co-stained cells. (L) Scatter boxplot quantifying cleaved caspase-3 positive cells per DG section analyzed. No significant changes were identified at E18.5 (P = 0.53; n=4), P7 (P = 0.28; n=4) or P14 (P = 0.11; n=4). Quantification of data presented as mean ± SEM. Scale bars = 50µm. Abbreviations: DG, dentate gyrus; gcl, granule cell layer; h, hilus; sgz, subgranular zone.

**Figure S3. *Mllt11* loss affected the formation of migrating Calbindin^+^ dentate granule cells.**

(A, B) Coronal sections of the developing DK at E18.5 in the WT and *Mllt11* cKO mouse stained for Calbindin (white) identifying nascent GCs. (C) Scatter boxplot quantifying a significant decrease in nascent GCs in the cKOs relative to WTs (P< 0.05, n=3). (D, E) Coronal sections of the dorsal hippocampus at P7 in the WT and cKO mouse stained for Calbindin. (F) Scatter boxplot identifying a significant increase in the formation of nascent GCs in the cKOs

## Contributions

SM sectioned, immunostained, analyzed the data, conducted microscopy generated figures, and wrote the first draft. DST assisted with sample preparation, data acquisition, microscopy and writing. KH conducted primary neural stem cell culture analysis and figure preparation. EAW assisted with data analysis and figure preparation. AI supervised the project, obtained funding, assisted with microscopy, edited the figures, co-wrote and edited the manuscript.

## Abbreviations

DG,: dentate gyrus;
GC,: granule cells;
GCL,: granule cell layer;
NP,: neural progenitor;
SGZ,: subgranular zone.

