## Supplementary figures and images for "Impaired subgranular zone radial glia morphology and transient amplification of neural progenitors in *Mllt11*-deficient mice leads to increased hippocampal neurogenesis"

### Supplemental Figure S1

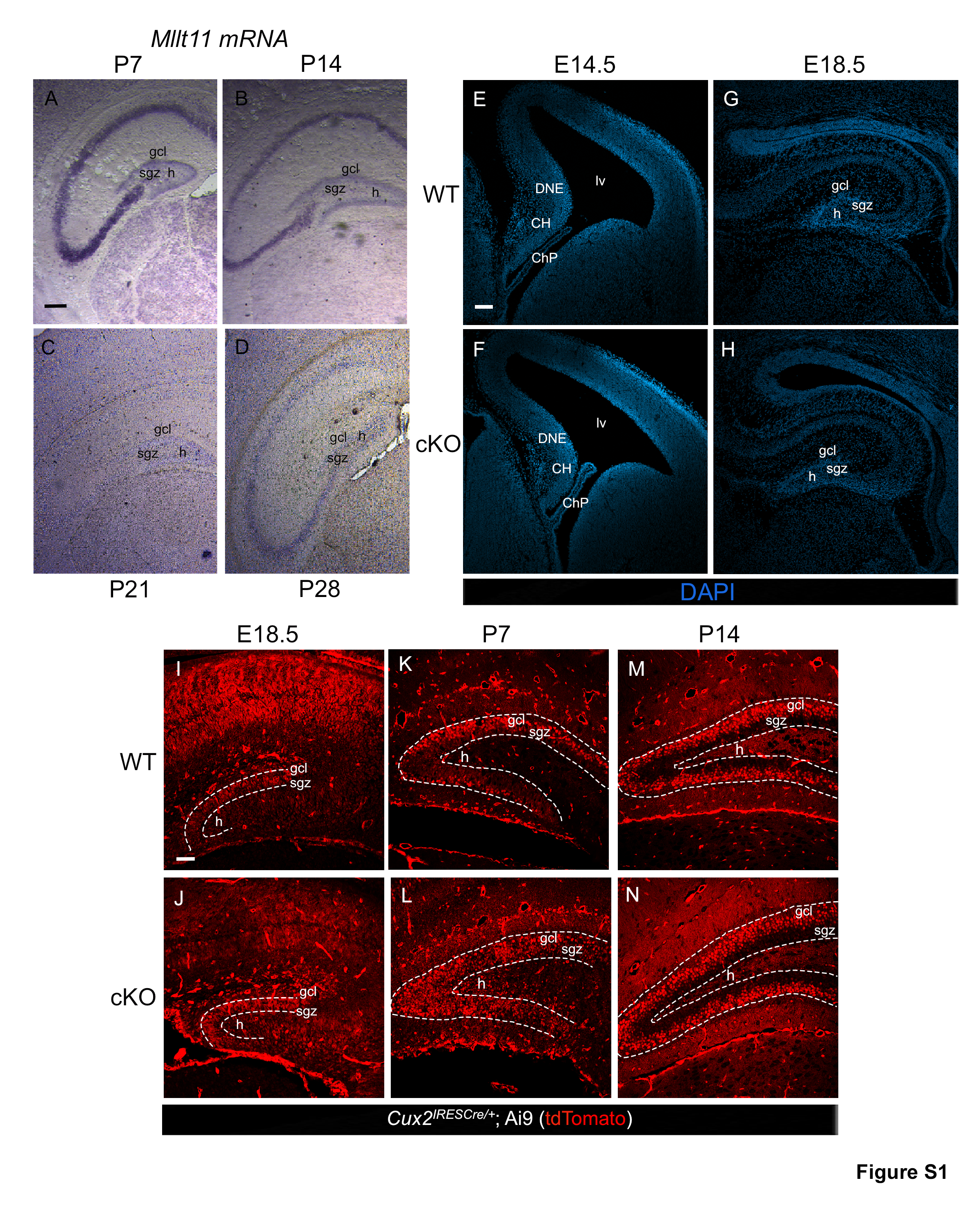

### Supplemental Figure S2

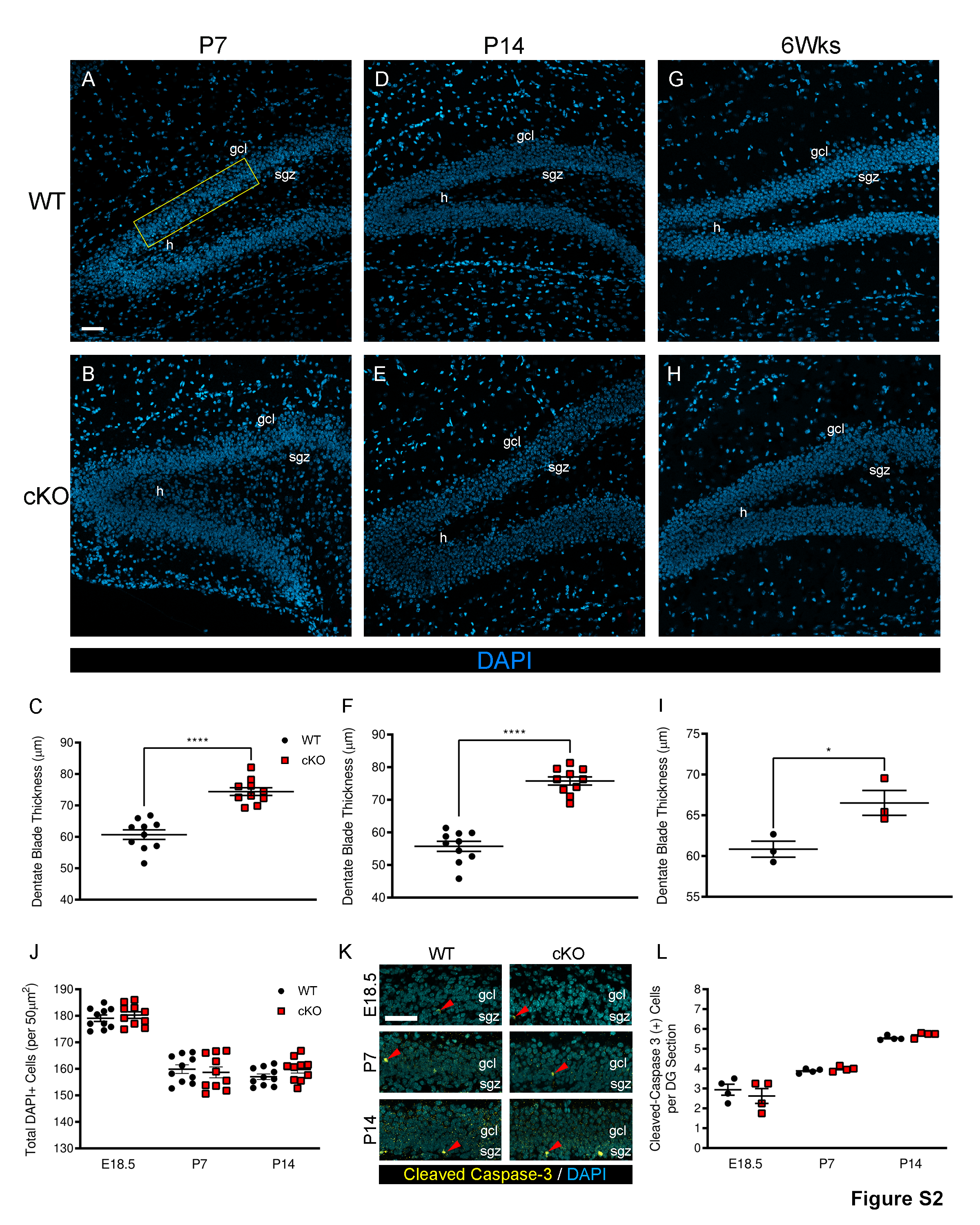

### Supplemental Figure S3

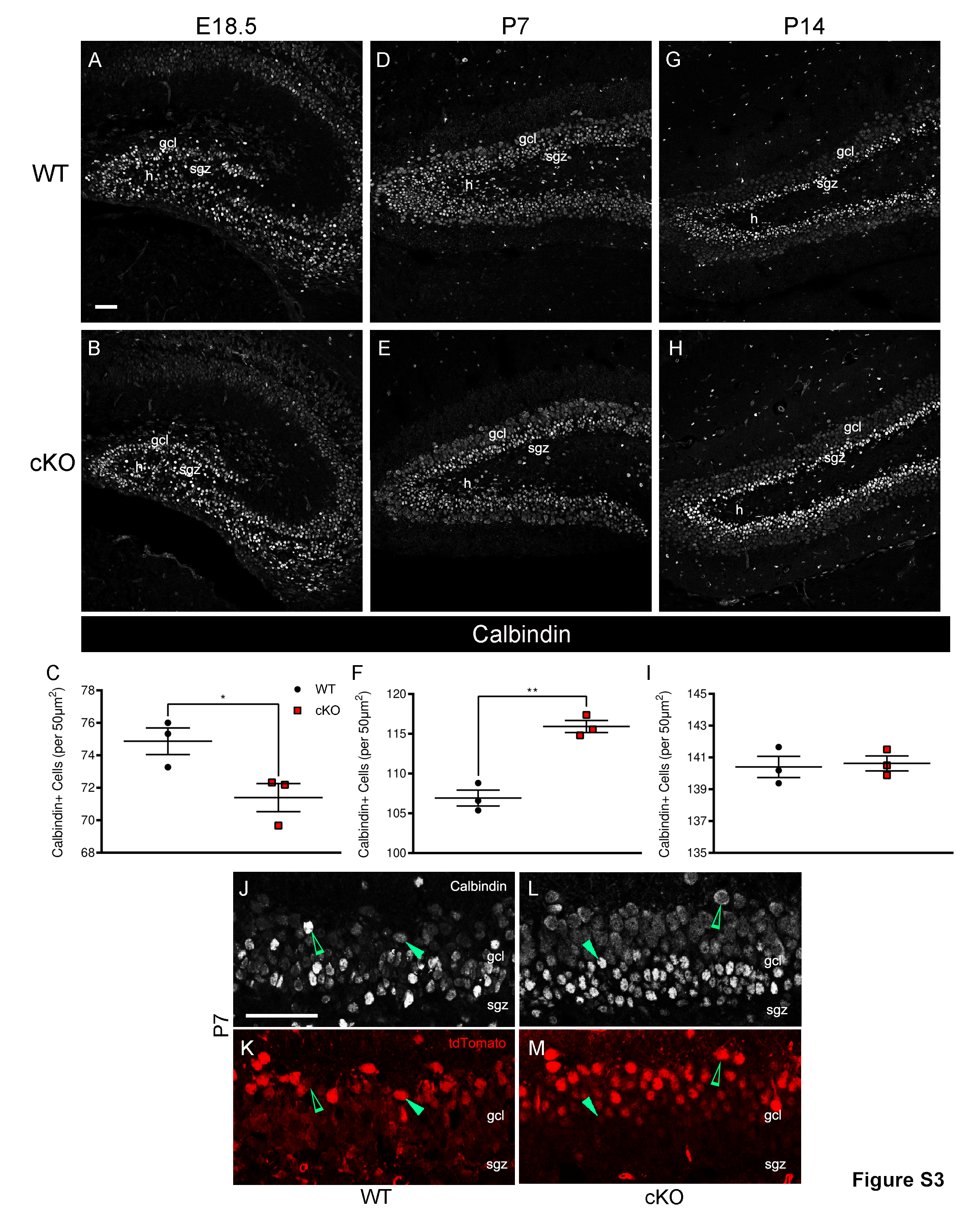
